# Seeing beyond the target: Leveraging off-target reads in targeted clinical tumor sequencing to identify prognostic biomarkers

**DOI:** 10.1101/2021.05.28.446240

**Authors:** Serghei Mangul, Jaqueline J Brito, Stefan Groha, Noah Zaitlen, Alexander Gusev

## Abstract

Clinical tumor sequencing is rapidly becoming a standard component of clinical care, providing essential information for selecting amongst treatment options and providing prognostic value. Here we develop a robust and scalable software platform (SBT: Seeing Beyond the Target) that mines discarded components of clinical sequences to produce estimates of a rich set of omics features including rDNA and mtDNA copy number, microbial species abundance, and T and B cell receptor sequences. We validate the accuracy of SBT via comparison to multimodal data from the TCGA and apply SBT to a tumor panel cohort of 2,920 lung adenocarcinomas to identify associations of clinical value. We replicated known associations of somatic events in *TP53* with changes in rDNA (p=0.012); as well as diversity of BCR and TCR repertoires with the biopsy site (p=2.5×10^−6^, p<10^−20^). We observed striking differences in EGFR mutant lung cancers versus wild-type, including higher rDNA copy number and lower immune repertoire diversity. Integrating clinical outcomes, we identified significant prognostic associations with overall survival, including SBT estimates of 5S rDNA (p=1.9×10^−4^, hazard ratio = 1.22) and TCR diversity (p=2.7×10^−3^, hazard ratio=1.77). Both novel survival associations replicated in 1,302 breast carcinoma and 1,651 colorectal cancer tumors. We anticipate that feature estimates derived by SBT will yield novel biomarker hypotheses and open research opportunities in existing and emerging clinical tumor sequencing cohorts.

## Introduction

Health care systems and private companies are now routinely sequencing tumors as part of clinical care in order to identify actionable mutations and improve patient care (DFCI OncoPanel^1^, MSK IMPACT^2^, MGH SNAPSHOT^3^, Foundation ONE ^4^, Tempus xT^5^). To reduce costs, tumor sequencing platforms only sequence exons from a small number of known cancer genes^1,2^. However, capture technologies are imperfect, and a substantial number of off-target reads are produced across various sequencing technologies^6,7^. We previously used off-target reads of clinical tumor sequences to impute genotypes and construct germline research cohorts^8^. Here, we extend this work with integrated multiple genomics methods^6,9^ to develop “Seeing Beyond the Target” (SBT), a software platform that mines discarded and off-target reads from existing tumor sequencing projects to produce a rich set of omics features. We show that SBT can uncover components of the tumor microenvironment that may serve as prognostic biomarkers.

We performed benchmarking using whole exome and whole transcriptome data to show that SBT can accurately estimate T and B cell receptor sequence diversity^10^; microbial, ribosomal^11^, and mitochondrial^12,13^ profiles. We demonstrated the utility of SBT via an analysis of 2,920 LUAD tumors with panel sequencing from the Dana-Farber *Profile* cohort^1,2^ — a prospective collection of patient biopsies -- replicating 6 published discoveries^6,11,12,14–16^ that previously required specifically designed experiments **(Table S1)**. We then leveraged the SBT inferred features to identify novel prognostic biomarkers, including estimates of 5S rDNA copy number and TCR α diversity, which were significantly associated with overall survival across multiple cancer types. The SBT implementation is highly efficient **(Figure S1**)^17^ and freely available, enabling similar utilization of the growing number of tumor-sequencing cohorts^1–5^.

## Results

### SBT is a comprehensive platform able to extract diverse omics features from targeted clinical tumor sequencing

Here we report the development of SBT, a computational platform able to extract a rich set of omics features directly from off-target tumor data. First, SBT estimates mtDNA and rDNA copy numbers estimated using reads mapping to the mtDNA genome and various rDNA repeat regions. Second, SBT uses ImReP^9^, our recently developed method to assemble T and B cell receptor clonotypes from off-target tumor reads. Common features of the immune repertoire inferred by ImReP include diversity and richness of various chains of BCR and TCR repertoires. Lastly, SBT uses reads mapped to microbial genomes to estimate microbial load. All features inferred by SBT are adjusted for off-target coverage. The full list of SBT features is provided in Table S4.

### Overview of datasets

We applied the SBT pipeline to four large-scale cancer sequencing cohorts: lung adenocarcinoma (LUAD) tumors from the TCGA with whole-exome (WXS) sequencing for 558 and RNA-seq for 490 samples (“TCGA-LUAD”); 2,920 LUAD cancer tumors (1,720 primary tumors) sequenced on the OncoPanel^1^ (“OP-LUAD”) used for discovery; as well as 1,302 primary breast carcinoma tumors (“OP-BRCA) and 1,651 primary colorectal cancer tumors (“OP-CRC”) sequenced on the OncoPanel and used for cross-cancer replication. The OncoPanel is a hybrid capture tumor sequencing platform targeting 200-500 cancer-related genes across three-panel versions^1^ (see Methods). The OncoPanel cohorts were prospectively collected as part of standard clinical care at the Dana-Farber Cancer Institute and consented for research.

### Inference of the immune repertoire yields novel associations with somatic drivers

We assayed T and B cell receptor sequences to characterize the immune microenvironment across patients diagnosed with non-small cell lung cancers. We have previously shown that ImReP using RNA-Seq can accurately estimate the relative abundance of BCR clonotypes and can capture a significant portion of the BCR repertoires captured by targeted BCR-Seq and other methods (Figure S5). Here, we validated the ImReP method for targeted tumor DNA sequencing.

Using 482 TCGA-LUAD samples sequencing by both RNA-Seq and WXS, we investigated the portion of the RNA-Seq-based immune repertoire captured by tumor WXS. ImReP using WXS data was able to capture more than 50% of the TCRB, TCRG, IGK, and IGL RNA-Seq-based clonotypes with a frequency higher than 10% and a smaller fraction of the repertoire of other chains (Figure S7). However, due to the decreased number of receptor-derived reads provided by WXS compared to RNA-Seq (Figure S6), the majority of WXS-based immune repertoire features were not associated with previously reported RNA-Seq-based measurements of the immune landscape^15^ (Figure 2). Features of T cell receptor beta chain estimated by ImRep from WXS data were Bonferroni significantly associated with infiltration levels of CD8+ T Cells (TCR β richness: effect=0.33, p=3.0×10^−4^). Additionally, features of immunoglobulin kappa chains estimated by ImReP based on WXS data were significantly associated with RNA-Seq-based overall BCR richness estimated as the total number of distinct immunoglobulin clonotypes (IG κ richness: effect=0.49, p=6.6×10^−4^).

We evaluated changes in the immune repertoire associated with recurrent somatic mutations in the OP-LUAD cohort (Figure 1). To ensure sufficient power, we restricted to three genes with highly recurrent mutations in at least 20% of the cohort: EGFR, KRAS, and TP53. We identified Bonferroni significant associations between somatic mutations in *EGFR* and BCR richness defined as the number of distinct clonotypes (effect=-0.099, p=9.1×10^−5^) and BCR infiltration measured based on the number of BCR-derived reads (effect=-0.099, p=6.8×10^−5^), potentially suggesting the presence of *EGFR*-specific antibody resistance^18^. Additionally, we detected novel associations between somatic mutations in *KRAS* and measures of the TCR repertoire (TCR γ richness: effect=0.062, p=3.0×10^−3^), which have not been previously observed^15,16^. Overall the increased sample size of our study enabled the detection of multiple associations between adaptive immune repertoires features and somatic driver alterations.

**Figure 1.**
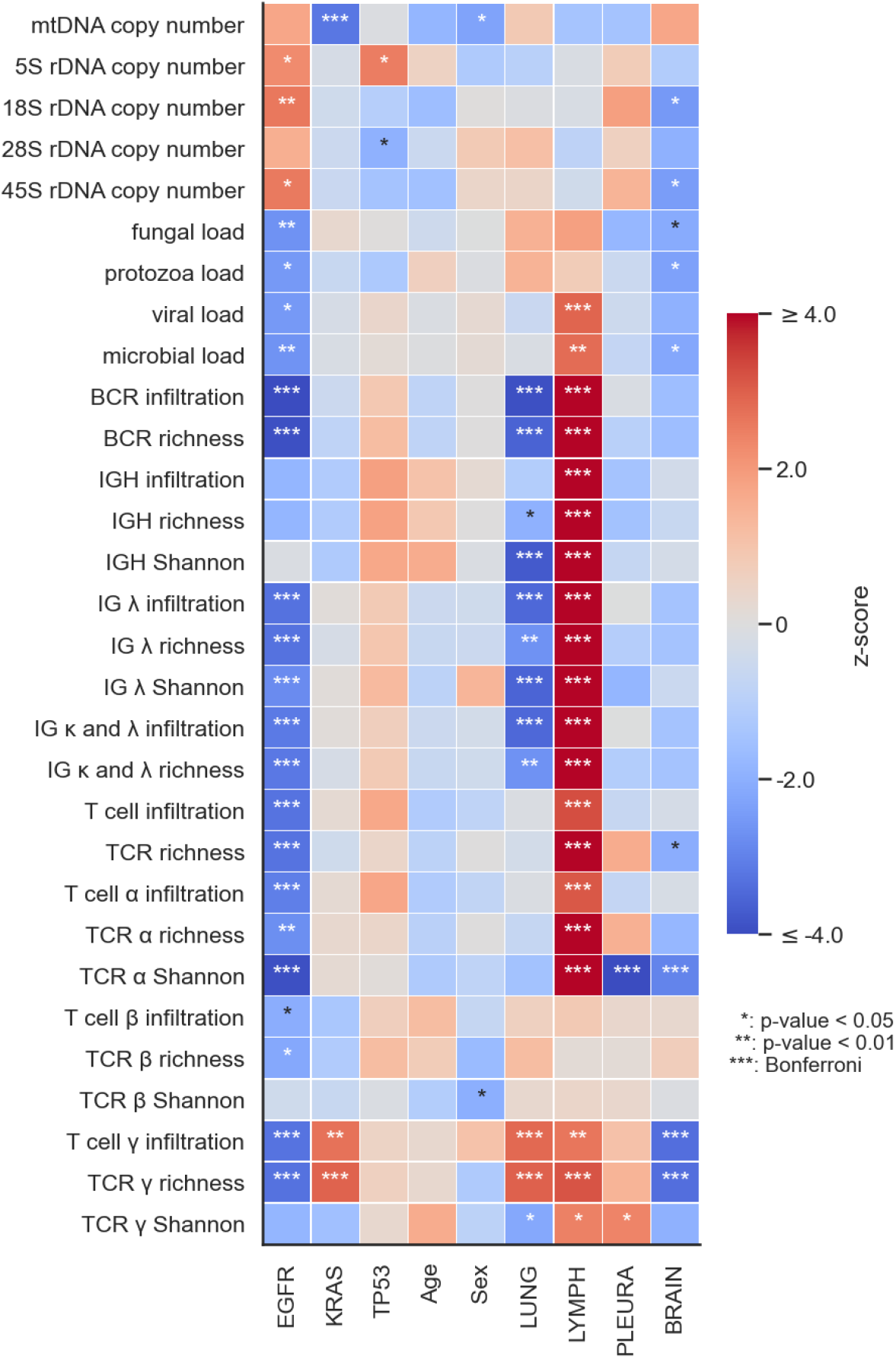
Association of SBT features with clinical factors and somatic mutations in the OP-LUAD cohort. Covariates are average off-target coverage, panel version, and tumor purity. Significant results are indicated with stars. Colors red and blue represent positive and negative association effect directions, respectively.

We used metastatic OP-LUAD samples biopsied from body sites with extensive immune activity (e.g. lymph nodes) as a positive control of the immunogenomics features inferred by ImReP. We observed highly significant associations between clonotype diversity of both BCR and TCR repertoires with the biopsy site, as has been previously detected using RNA-Seq data^14,16^. For example, we detected the association between TCR α Shannon and lymph vs non-lymph (effect=0.12, p=2.5×10^−6^), and IGH Shannon and lymph vs non-lymph (effect=0.26, p-value<10^−20^) (Figure 1). Restricting to just the lung biopsied tumors, we additionally observed significant associations between richness of immunoglobulin heavy chain repertoire and TMB (effect=0.26, p=9.1×10^−3^) that were not observed when including tumors from other sites (Figure S3).

### Inference of mtDNA abundance yields associations with mutations in KRAS

We next investigated the ability of tumor-only sequencing to estimate mitochondrial DNA (mtDNA) copy number variation. The involvement of mitochondria in essential functions such as biosynthesis, signaling, cellular differentiation, apoptosis, and cell growth can play a critical role in tumorigenesis^19,20^. WGS provides a unique opportunity for a comprehensive molecular characterization of mitochondria across a broad range of cancer types^21^, but remains cost-prohibitive in clinical cohorts^22^. Previously, it has been shown that targeted DNA sequencing data can provide robust mtDNA copy number estimates comparable to one derived from WGS data^11,12^.

For each cohort, we remapped DNA reads to the mitochondrial genome with consistent settings to ensure consistent processing of mitochondrial reads (see Methods). Using the TCGA-LUAD data, we showed that SBT mtDNA copy number estimates, calculated as a ratio between mtDNA coverage and average coverage of off-target regions, were correlated with previous estimates computed as the ratio of mtDNA reads to nuclear DNA reads (Pearson correlation=0.61, p-value<10^−20^) (Figure S2)^12^, demonstrating consistency with previous work. It was previously shown that mtDNA abundance can be correlated with somatic mutations in key oncogenes^12^. For example, increased mtDNA abundance was determined to be associated with mutations in *TP53* in endometrial cancer^12^. Here we investigated associations between mtDNA abundance and somatic mutations in the OP-LUAD cohort. We again restricted to the highly recurrent putative driver genes *EGFR, KRAS*, and *TP53*. We observed a Bonferroni significant negative association between mtDNA abundance and somatic mutations in *KRAS* (effect=-0.046, p=1.3×10^−3^) (Figure 1). This association with *KRAS* remained significant after restricting to just samples biopsied from the lung (effect -0.051, p=6.6×10^−3^; Figure S3). We additionally observed nominal associations between mtDNA abundances and sex (Figure 1, Figure S3).

### Inference of rDNA abundance recapitulates associations with mutations in TP53

We next investigated the ability of tumor-only sequencing to estimate ribosomal DNA (rDNA). Human ribosomes are regulators of translation and play an essential role in tumorigenesis^23^. Ribosomal defects and perturbation can have a direct effect on tumor progression^24^. We have developed an approach with low computational requirements able to effectively leverage originally mapped reads and extract the candidate rDNA reads which are further carefully mapped to rDNA complete repeating units (see Methods). After applying this approach here, we computed rDNA copy number estimates as the ratio between the coverage of reads mapped to individual rDNA components and average coverage of off-target regions.

As with mtDNA, we evaluated the association of rDNA abundances with recurrent somatic mutations in the OP-LUAD cohort. We observed a high correlation between rDNA copy number estimates of individual rDNA components (e.g. 28S vs 18S: Pearson correlation=0.87, p-value<10^−20^; Figure S4), consistent with previous findings^11^. We confirmed the known associations of somatic events in *TP53* with changes in the rDNA components (5S rDNA: effect=0.041, p=0.012; 28S rDNA: effect=-0.034, p=0.050)^11^ (Figure 1), which also replicated in TCGA-LUAD WXS samples (e.g., 18S rDNA: effect=-0.066, p=5.4×10^−3^) (Figure 2). In addition, we observed significant novel associations between mutations in *EGFR* and changes in the rDNA components (e.g., 18S rDNA: effect=0.037, p=9.4×10^−3^) (Figure 2).

**Figure 2.**
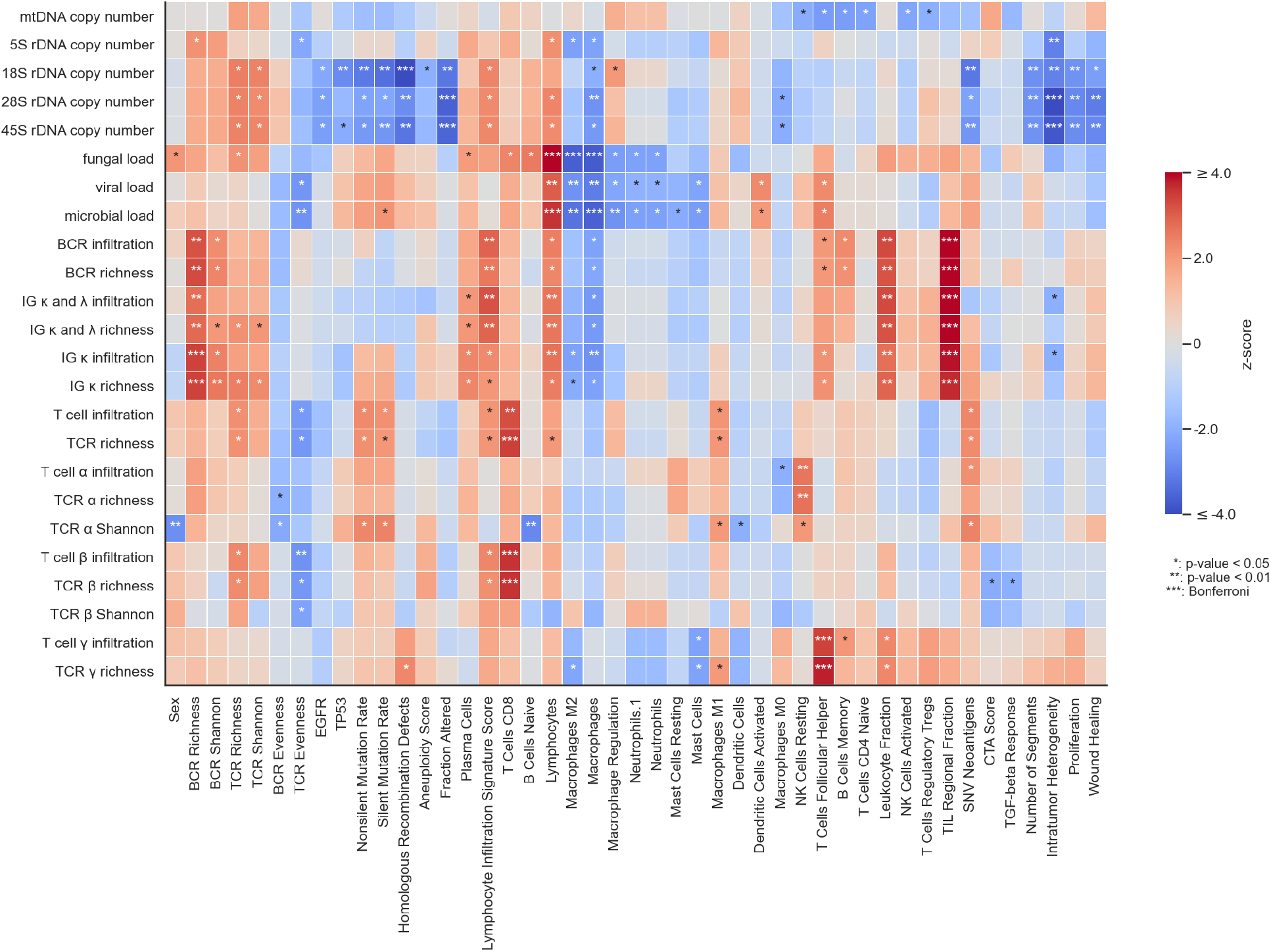
Association of SBT features with clinical factors and measurements from the immune landscape^15^ for TCGA-LUAD WXS samples. Covariates are the average off-target coverage and tumor purity. Significant results are indicated with stars. SBT features with no significant results are removed. Colors red and blue represent positive and negative association effect directions, respectively.

## Inference of microbial load yields novel associations with mutations in EGFR

There is increasing evidence of the importance of intratumor microbiome across various tumor types^25–27^ and tumor type-specific intracellular microbiomes have been described across various tissue and cancer types^28^. We and others have previously demonstrated that reads of microbial origin often occur in sufficient abundances among human reads to access microbial loads and composition^6,10,22,25^. SBT uses reads aligned to the viral, protozoa, and fungi genomes to access the microbial loads across patients. Comparing the RNA-seq and WXS data, we observed a significant correlation for the overall microbial load, but not individual loads (Table S2). This may suggest a distinct ability of these different sequencing protocols to capture individual microbial components. Applying to the OP-LUAD data, we observed a significant negative association of microbial loads with somatic mutations in *EGFR* (e.g., fungal load: effect=-0.034, p=7.8×10^−3^) (Figure 1). Similar to our previous analysis in GTEx tissues^6^, we observed substantial microbial loads across tumor biopsy sites (Figure S8) with all loads nominally lower in biopsies from brain versus non-brain (Figure 1).

### SBT features can serve as prognostic biomarkers across cancers

We next sought to identify SBT features associated with overall survival in 1,720 OP-LUAD primary tumors. We applied Cox proportional-hazard models, with sequencing date taken as the start point and death (loss to follow-up) taken as the event (censoring) points. Age, sex, tumor purity, and technical features were included as covariates. Features violating proportional hazards assumption were removed (see Methods).

The SBT features yielded 9 significant associations (q-value <0.1; Figure 3), with the most significant associations (q-value <0.05) found for 5S rDNA copy number (q-value=7.6×10^−3^, hazard ratio (HR) = 1.22) and TCR α Shannon (q-value=5.2×10^−2^, HR=1.77). Other significant associations (q-value <0.1) were observed for viral load, microbial load, protozoa load, TCR α richness, TCR γ richness, TCR β richness, and T cell β infiltration. Of these associations, 5/9 remained significant (q-value <0.1) and had a consistent effect when tested in OP-LUAD non-primary tumors (n=1,200 recurrent or metastatic tumors), serving as replication in held-out samples.

**Figure 3.**
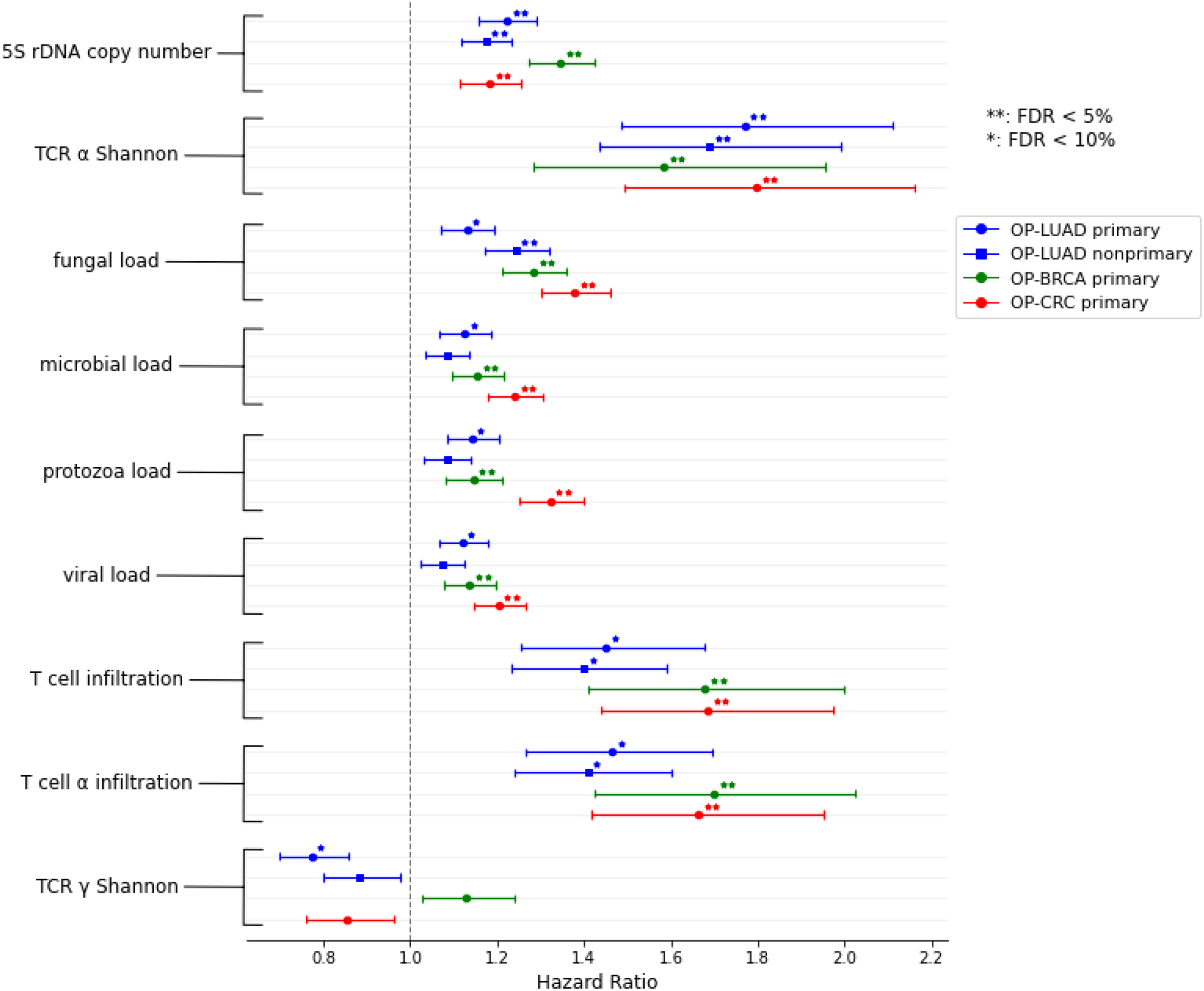
Hazard ratio from significant associations of SBT features with the survival time obtained from the Cox proportional-hazard model applied over the OP-LUAD cohort (n=1,720 primary tumors and 1,200 nonprimary tumors) and used for discovery, and breast carcinoma (only female, 1,302 primary tumors) and colorectal cancer tumors (n=1,651 primary tumors) sequenced on the OncoPanel and used for cross-cancer replication.

To further replicate the associations observed for LUAD samples, we analyzed two additional cancer types from the same sequencing cohort: primary breast carcinoma tumors (n=1302) and primary colorectal cancer tumors (n=1651). Of the 9 significant (q-value<0.1) associations in OP-LUAD, 8/9 replicated in breast cancer and 8/9 replicated in colorectal carcinoma (Figure 3). Notably, we observed highly significant associations in both cohorts with 5S rDNA copy number (breast: p=2.2×10^−7^, HR=1.35; colorectal: p=4.6×10^−3^, HR=1.18) and TCR α Shannon (breast: p=8.6×10^−3^, HR=1.58; colorectal: p=7.3×10^−3^, HR=1.80). Kaplan-Meier curves (which do not account for covariates) showed a clear separation of median survival time and proportionality (Figure 4).

**Figure 4.**
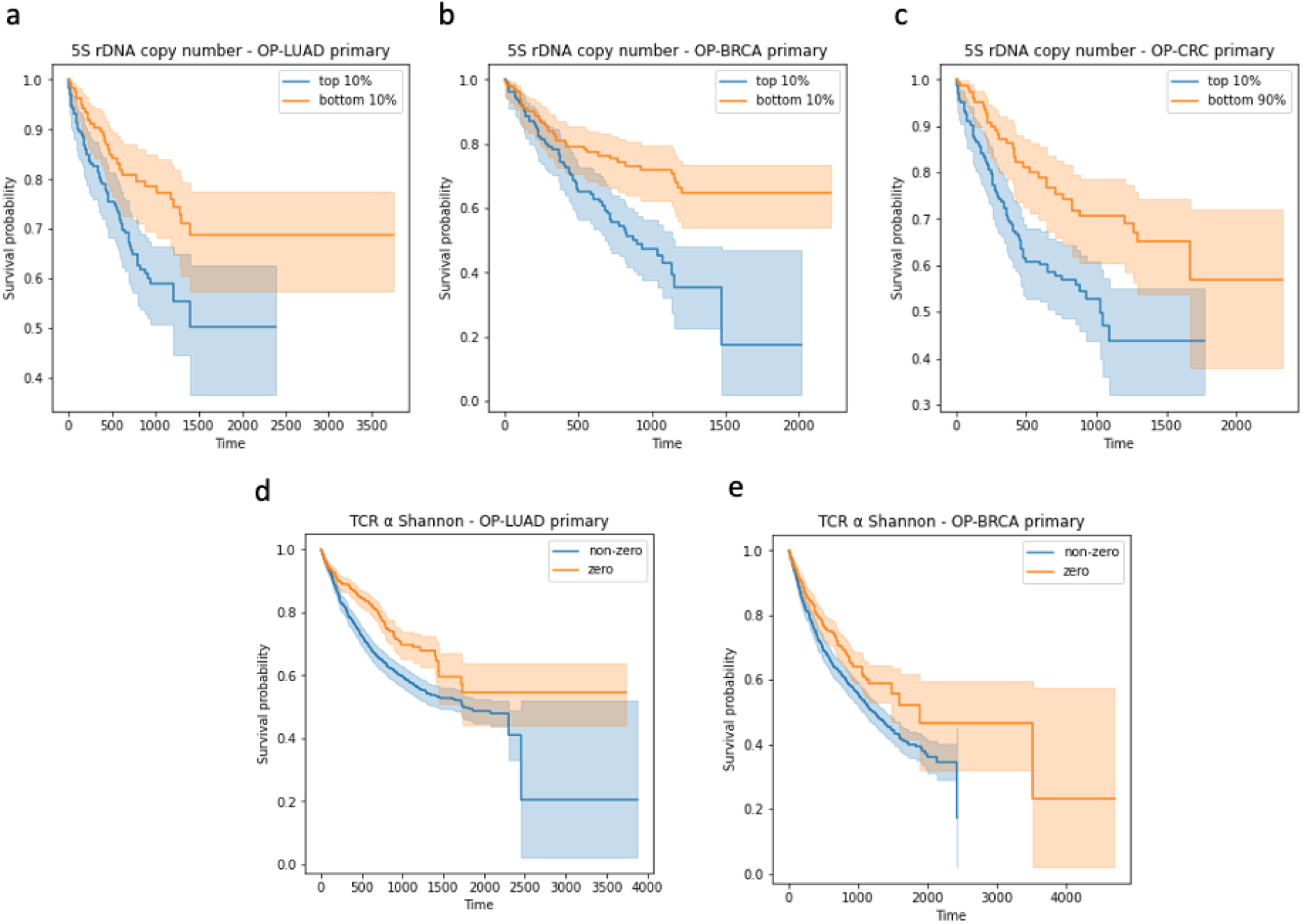
Kaplan-Meier survival plots displaying the top and bottom 10% values of 5S rDNA copy number and zero and non-zero values of TCR Shannon of primary tumors: a) 5S rDNA copy number (OP-LUAD primary); b) 5S rDNA copy number (OP-BRCA primary); c) 5S rDNA copy number (OP-CRC primary); d) TCR α Shannon (OP-LUAD primary); and e) TCR α Shannon (OP-BRCA primary).

### Associations between domains of SBT features

To lend further context to the features inferred by SBT from DNA, we turned to the TCGA-LUAD samples for which the immune landscape had recently been characterized using multiple molecular techniques^15^ (Figure 2). 5S rDNA copy number, which we found to be associated with poor survival (see above), was positively associated with the fraction of lymphocytes in TCGA data (Figure 2). These results are consistent with previous findings that lymphocyte infiltration levels are associated with poor survival^15^ which may be indirectly captured by SBT features.

More broadly, we observed widespread associations between SBT features and TCGA-LUAD immune features. Bonferroni significant (and negative) associations were observed between rDNA features and broad tumor instability: including 18S and Homologous Recombination Deficiency (effect=-0.19, p=6.5×10^−5^); 28S and Fraction Altered (effect=-0.15, p=3.1×10^−4^); 28S and Intratumor Heterogeneity (effect=-0.18, p=3.9×10^−5^); with similar patterns for 45S. Bonferroni significant associations were also observed between fungal/microbial load and increased Lymphocyte counts (p=1.9×10^−5^) and decreased Macrophage counts (p=2.0×10^−4^). Lastly, Bonferroni significant associations were observed between (WXS) IG κ infiltration and (RNA) BCR richness; as well as (WXS) TCR richness and (RNA) CD8 T Cell counts. In summary, SBT applied to WXS captured broad components of tumor instability, infiltration, and TCR/BCR expansion without the cost of additional sample collection and sequencing.

## Discussion

Cancer sequencing studies, like TCGA^29^, ICGC^30^, and POG570^15,22^ have provided unprecedented functional genomic resources for cancer research. However, sample sizes are small relative to genetic studies, much of the clinical data are sparse or outdated^31^, and the patient populations have limited longitudinal follow-up. Targeted cancer sequencing, an increasingly routine component of clinical care (e.g. AACR GENIE^32^), offers reduced costs and rich clinical data collected across hundreds of thousands of cancer patients but is limited in genomic scope. To overcome this limitation, we have developed the SBT computational platform, able to extract multiple omics measurements directly from off-target tumor data. To illustrate the power of leveraging off-target reads in targeted clinical tumor sequencing we applied SBT to a tumor panel cohort of 2,920 lung adenocarcinomas, identifying both known and novel associations with recurrent somatic drivers, as well as prognostic biomarkers for survival that replicated across multiple cancer types. Previous studies attempting to determine associations of various omics features with survival outcomes were typically underpowered due to the small sample size of each cancer type^11,16^, and recent work detected an association of increased TCR diversity with a progression-free interval in various cancer types^15^, but not LUAD.

It is important to note that there are several shortcomings of using omics features inferred from targeted tumor sequencing. First, in contrast to RNA-Seq data where cell type composition can be inferred computationally based on gene expression^33^, targeted sequencing lacks the capability to infer the cell-type composition of the sample and adjust for cell type confounding. A potential solution is to utilize the reads derived from T and B cell receptors to estimate the fraction of T and B cells. However such methods need to be carefully validated based on comprehensive gold standard data. Second, the proposed methods and their accuracy might be platform-specific and downstream analyses must account for platform-specific biases. Additionally, it is challenging to accurately capture many omics features due to the variability of sequencing data quality. For example, it has been recently shown that many reported ribosomal copy number estimates may be an artifact of sequencing data quality and not reflect real biological signals^34^, which may apply to SBT estimates. Finally, methods for various SBT features remain in active development, and the proposed software platform must be updated to keep up with best practices. For example, estimating components of the microbiome from off-target reads is especially difficult, and better methods are continually being developed^35^.

Additional caveats apply to our analyses of patient survival. The clinical cohort analyzed here is a heterogeneous population and, while panel sequencing is not uncommon, patients are still often sequenced due to having an advanced disease or poor response to therapy and may not be reflective of the broader patient population. Data on the date of diagnosis, stage at diagnosis, and detailed cancer subtypes (e.g. hormone receptor status for breast tumors) were generally unavailable and could not be incorporated into the analysis. While the prognostic SBT features replicated across diverse cancer types and thus likely reflect associations in this population, they should not be interpreted as causal. In TCGA data, SBT features were correlated with many diverse measures of the tumor and tumor microenvironment, and in the clinical data they were associated with multiple key somatic events including EGFR driver status and TMB. Prognostic SBT features are thus likely to be only surrogates for underlying causal mechanisms that remain to be identified. We envision that our analysis framework can be used to efficiently identify putative prognostic features across thousands of patients, which can then be investigated in detail and validated with richer, multi-modal studies.

Despite these shortcomings, we anticipate SBT will increase the scientific value of many existing cancer cohorts. To further the ability of other groups to conduct similar research we have released SBT on the cloud and as a fully free and downloadable software package at https://github.com/Mangul-Lab-USC/sbt.

## Methods

### OncoPanel data collection and processing

OncoPanel samples were collected as part of the Dana-Farber Profile prospective tumor sequencing cohort^36^. Patients were recruited based on available material and consent and were not otherwise ascertained for age, sex, stage, or tumor site. All patients provided informed consent for research (Institutional Review Board (IRB) protocols 11-104 and 17-000) and secondary analysis of data was approved by the IRB (protocol 19-033). The OncoPanel platform is a next-generation sequencing assay that targets the exons of 275-447 cancer genes on one of three panel versions. Sequences were aligned to the human genome (hg19) using bwa and processed with the GATK IndelRealigner. Somatic mutations were called using MuTect (v1.1.4) relative to a de-identified panel of normal samples. The sequencing and variant calling pipeline was used for clinical reporting to patients and physicians and has previously been validated to achieve 98% sensitivity and 100% specificity for single nucleotide variants^1^.

### Datasets used for SBT validation

To investigate the accuracy of SBT, we first applied it to a variety of high-quality data sets with various types of existing tumor and normal sequencing, genotyping, and RNA-sequencing data (**Table S3**). This analysis included: 2,920 lung adenocarcinoma cancer tumors sequenced on the OncoPanel (“OP-LUAD”) and used for discovery, and 2,384 breast carcinoma (only female) and 2,278 colorectal cancer tumors sequenced on the OncoPanel and used for replication. We also analyzed tumor whole-exome sequencing (WXS) in TCGA: 558 WXS and 490 RNA-seq LUAD samples (“TCGA-LUAD”). TCGA-LUAD is composed of sequencing data of cancer patients and controls were downloaded from GDC data portal (https://portal.gdc.cancer.gov/). We have downloaded individuals with matching (482 samples) whole exome (WXS) RNA Sequencing (RNA-Seq) samples. Data was downloaded as a BAM file with mapped and unmapped reads. Both WXS and RNA-Seq reads were mapped to hg38 reference using a bwa aligner. Parameters, input, and output data are shared for publicly available data sets (https://github.com/Mangul-Lab-USC/sbt_publication). For each data type output, we examined multiple metrics against an accepted gold standard, which we describe in turn below.

### Preparing input reads for SBT platform

References used to map reads contain multiple non-human references, including HPV, HBV, and HCV viruses. We have added reads mapped to those references to the unmapped reads (extended unmapped reads). SBT platform is a natural extension of our previously developed method (ROP, https://github.com/smangul1/rop), adjusted for the targeted cancer data including WXS and OncoPanel data able to infer various omics features. The definitions of SBT features are detailed in Table S4.

### Estimation of off-target reads coverage

To account for the total number of reads of the sample and the efficiency of the caption, we used the per-base depth of the intergenic regions. Intergenic regions are not captured by any modern targeted sequencing protocols, such as Whole Exome Sequencing (WXS) and OncoPanel (OP), and can be used as an effective method to estimate background reads depth. Reads mapped to intergenic regions are extracted from the BAM file using samtools^37^. Coverage of intergenic regions is generated using the samtools depth function. The off-target coverage is computed as the sum of the depth of the intergenic regions divided by the sum of the size of these regions.

### Estimation of rDNA copy number

We downloaded the human 5S repeat region of length 2231 bp (GenBank ID X12811.1), 18S (GenBank ID X03205.1), and 28S (GenBank ID M11167.1) units composing Ribosomal DNA complete repeating unit. We refer to those individual references as the rDNA references. A straightforward approach is to align original raw reads to the rDNA references. However, this approach is computationally intractable given the increasing number of reads being generated by the sequencing protocols. Considering only unmapped reads can be risky due to homology between rDNA sequence and the human reference genome. For example, it was previously demonstrated that reads are simultaneously mapped to GL000220.1 scaffold and ribosomal DNA complete repeating unit. Similarly, reads mapped to 5S repeat regions are also mapped to 1q42 (hg19, chr1:226743523–231781906). We used a comprehensive method to identify regions of the human genome that are homologous to ribosomal DNA sequences. We simulated 75bp overlapping substring from ribosomal DNA sequences (kmers) and detected regions where rDNA substring are mapped. Reads mapped to these regions are extracted from the BAM file using samtools^37^ and added to the unmapped reads. The extended set of unmapped reads are mapped to rDNA references using Bowtie2 aligners (v2.3.5.1). Pairing information was disregarded and reads were aligned in end-to-end mode (–end-to-end). rDNA reads were stored in a BAM file. Coverage of rDNA regions is generated using the samtools depth function.

### Estimation of mtDNA copy number

We downloaded the human mitochondrial genome (NCBI ID NC_012920.1). We extracted reads from the original BAM files, which are mapped to the mitochondrial genome. Resulted reads were mapped to mtDNA reference using bowtie2 in end-to-end mode (–end-to-end), similarly as was done for rDNA copy number estimation.

### Characterization of T and B cell receptor repertoires

We used ImReP (https://github.com/Mangul-Lab-USC/imrep) to extract receptor-derived reads and assemble T and B cell receptor clonotypes. We run ImReP over the BAM files using the options “noCast” and “noOverlapStep”.

### Estimation of microbial load

We have mapped prepared input reads (See Section Prepare input reads for SBT platform) onto the collection of viral, fungi, and protozoa reference genomes downloaded from NCBI (ftp://ftp.ncbi.nih.gov/). We used bwa mem with default settings. Similarly, to ROP (https://github.com/smangul1/rop) we considered reads mapped with 90% identity. Hits shorter than 80% of the input read sequence were removed (corresponding to 80bp of the 100bp read).

### Feature processing for association analysis

We applied an inverse normal transformation^38^ to covariates and outcome variables to account for their typically non-Gaussian distributions. In the OP-LUAD we normalized TMB; and in TCGA, we normalized all continuous outcomes variables. SBT features were often zero-inflated and required additional processing (Table S5). For SBT features with portions of zero values >10%, we created two variables: the first variable was dichotomous, indicating whether the SBT feature values was zero (set to 0) or non-zero (set to 1); the second variable was computed by applying the inverse normal transformation to the non-zero values and zero for the rest (this variable can be interpreted as an interaction between the first variable and the rank normalized feature). We discarded SBT features that had less than 30 non-zero values. For SBT features with portions of zeros < 10%, we applied the inverse normal transformation to all values.

### Association analysis

We use a linear regression model (statsmodels Python package) to test for association between SBT features and other genetic/immune features together with technical covariates. In OP-LUAD, technical covariates included tumor purity and off-target coverage; in TCGA WXS data, technical covariates included tumor purity and off-target coverage; in TCGA RNA data, technical covariates included tumor purity and the number of reads. For SBT features with portions of zeros <10%, we used a linear regression model with the inverse-normal transformed variable. For zero-inflated features that were divided into two variables as part of preprocessing (see above), we evaluated significance using a two-degree of freedom test. We performed two nested regressions: the first regression included both variables (and technical covariates) and the second only included the covariates. We then applied a two-degree of freedom likelihood ratio test to compute the p-value. The effect size to report in figures was determined by the highest absolute coefficient from the two variables.

### Survival analysis

For the survival analysis, we used the date of sequencing as the start date, death as the end-point (linked to the National Death Index), and censoring on the NDI linkage or clinical loss to follow-up. Taking sequencing as the start point eliminated potential immortal time bias (because all patients had to be sequenced to enter the cohort) and was robust to incomplete data, as the majority of patients were first diagnosed at other institutions. However, hazard ratios should be interpreted relative to this clinical reporting point rather than cancer diagnosis. For testing, we use a multivariate Cox proportional hazards model (lifelines Python package), controlling for age at sequencing (stratified into three equal groups), sex, biopsy site type (primary/non-primary), line of treatment, and tumor purity. We furthermore controlled for technical covariates: sequencing panel version and off-target coverage. For any covariate that violated the proportional hazards assumption at p<0.05/N (where N is the number of covariates), we stratified over the respective covariate. Any SBT features that violated the proportional hazards test were dropped. For each dataset, we applied a false discovery rate (FDR) correction to correct for multiple testing.

## Supporting information

Supplementary Data: Summary of results

## Acknowledgments

The authors would also like to acknowledge the DFCI Oncology Data Retrieval System (OncDRS) for the aggregation, management, and delivery of the clinical and operational research data used in this project; as well as the DFCI/BWH Data Sharing Group for the aggregation, management, and delivery of the genomics data used in this project.

## Data availability

All RNA-Seq and WES data discussed in this paper is available as part of The Cancer Genome Atlas (TCGA) Project. All data required to produce the figures and analysis performed in this paper are freely available at https://github.com/Mangul-Lab-USC/sbt-publication.

## Supplementary Data

Full summary statistics are provided in the Supplementary Data object.

## Code availability

SBT is freely available at https://github.com/Mangul-Lab-USC/sbt. SBT is distributed under the terms of the General Public License version 3.0 (GPLv3). All the code used to produce the results of this study, including figures, is available at https://github.com/Mangul-Lab-USC/sbt-publication.

## Supplementary Tables

**Table S1.** Summary of replicated results from previous studies.

| # | Summary of replicated results | Citation |
| --- | --- | --- |
| 1 | We observed highly significant associations between clonotype diversity of both BCR and TCR repertoires with the biopsy site, as has been previously detected using RNA-Seq data<sup>14,16</sup>. | <sup>14,16</sup> |
| 2 | Using the TCGA-LUAD data, we showed that SBT mtDNA copy number estimates, calculated as a ratio between mtDNA coverage and average coverage of off-target regions, were correlated with previous estimates computed as the ratio of mtDNA reads to nuclear DNA reads (Pearson correlation=0.61, p-value<10<sup>-20</sup>) (Figure S2)<sup>12</sup>, demonstrating consistency with previous work. | <sup>12</sup> |
| 3 | We observed a high correlation between rDNA copy number estimates of individual rDNA components (e.g. 28S vs 18S: Pearson correlation=0.87, p-value<10<sup>-20</sup>) in the OP-LUAD cohort (Figure S4), consistent with previous findings<sup>11</sup>. | <sup>11</sup> |
| 4 | We confirmed the known associations of somatic events in *TP53* with changes in the rDNA components (5S rDNA: effect=0.041, p=0.012; 28S | <sup>11</sup> |
|  | rDNA: effect=-0.034, p=0.050)<sup>11</sup> (Figure 1), which also replicated in TCGA-LUAD WXS samples (e.g., 18S rDNA: effect=-0.066, p=5.4x10<sup>-3</sup>) (Figure 2). |  |
| 5 | 5S rDNA copy number, which we found to be associated with poor survival (see above), was positively associated with the fraction of lymphocytes in TCGA data (Figure 2). These results are consistent with previous findings that lymphocyte infiltration levels are associated with poor survival<sup>15</sup> which may be indirectly captured by SBT features. | <sup>15</sup> |
| 6 | Similar to our previous analysis in GTEx tissues<sup>6</sup>, we observed substantial microbial load across various human tissues over the OP-LUAD cohort (Figure S8) with all loads nominally lower in biopsies from brain versus non-brain (Figure 1). | <sup>6</sup> |

**Table S2.** Pearson correlation of the microbial loads of RNA-Seq and WXS samples (n=482) from the TCGA-LUAD cohort.

| load | Pearson correlation | p-value |
| --- | --- | --- |
| fungal | 0.0028 | 0.95 |
| protozoa | -0.025 | 0.58 |
| viral | 0.074 | 0.10 |
| microbial | 0.29 | $< 10^{-20}$ |

**Table S3.** Overview of high-quality data sets. The number of samples in each dataset is presented (“n”). We documented the types of cancer each data contains (“Cancer Type”). We recorded the types of the sequencing technology used for samples in each study (“Tumor”). We documented whether samples from the study were sequenced by RNA-Seq (“RNA”). Abbreviations: WXS: Whole-Exome sequencing; OP: Oncopanel-sequencing; LUAD: Lung adenocarcinoma; BRCA: Breast carcinoma; CRC: Colorectal cancer.

| **Data Source** | **Cancer Type** | **Tumor** | **RNA** |
| --- | --- | --- | --- |
| OP-LUAD | Lung adenocarcinoma | OncoPanel (n=1,720<br>primary and<br>1,200 non-primary tumors) | - |
| OP-BRCA | Breast carcinoma | OncoPanel (n=1,302<br>primary tumors) | - |
| OP-CRC | Colorectal cancer | OncoPanel (n=1,651<br>primary tumors) | - |
| TCGA-LUAD | Lung adenocarcinoma | WXS (n=558) | RNA-Seq (n=490) |

**Table S4.** Definition of SBT features.

| Type | Feature | How it is measured |
| --- | --- | --- |
| mtDNA | mtDNA copy number | Measured based on reads covering mtDNA genome adjusted for off-target coverage |
| rDNA | 5S rDNA copy number | Measured based on reads covering 5S rDNA region adjusted for off-target coverage |
|  | 18S rDNA copy number | Measured based on reads covering 18S rDNA region adjusted for off-target coverage |
|  | 28S rDNA copy number | Measured based on reads covering 28S rDNA genome adjusted for off-target coverage |
| microbiome | fungal load | Measured based on number of reads mapped to fungal genomes adjusted for off-target coverage |
|  | microbial load | Measured based on number of reads mapped to microbial genomes adjusted for off-target coverage |
|  | protozoa load | Measured based on number of reads mapped to protozoan genomes adjusted for off-target coverage |
|  | viral load | Measured based on number of reads mapped to viral genomes adjusted for off-target coverage |
| Immune | IG $\kappa$ and $\lambda$ infiltration | Measured based on number of IGκ- and IGλ-derived reads adjusted for off-target coverage |
|  | BCR infiltration | Measured based on number of IG-derived (including both light and heavy chains) reads adjusted for off-target coverage |
|  | IGH infiltration | Measured based on number of IGH-derived reads adjusted for off-target coverage |
| | IG $\kappa$ infiltration | Measured based on number of IGκ-derived reads adjusted for off-target coverage |
| | IG $\lambda$ infiltration | Measured based on number of IGλ-derived reads adjusted for off-target coverage |
|  | T cell infiltration | Measured based on number of TCR-derived reads adjusted for off-target coverage |
| | T cell $\alpha$ infiltration | Measured based on number of TCRα-derived reads adjusted for off-target coverage |
| | T cell $\beta$ infiltration | Measured based on number of TCRβ-derived reads adjusted for off-target coverage |
| | T cell $\delta$ infiltration | Measured based on number of TCRδ-derived reads adjusted for off-target coverage |
| | T cell $\gamma$ infiltration | Measured based on number of TCRγ-derived reads adjusted for off-target coverage |
| | IG $\kappa$ and $\lambda$ richness | Measured based on number of distinct IGκ and IGλ clonotypes adjusted for off-target coverage |
|  | BCR richness | Measured based on the number of distinct IG (including both light and heavy chains) clonotypes adjusted for off-target coverage |
|  | IGH richness | Measured based on the number of distinct IGH clonotypes adjusted for off-target coverage |
| | IG $\kappa$ richness | Measured based on the number of distinct IGκ clonotypes adjusted for off-target coverage |
| | IG $\lambda$ richness | Measured based on the number of distinct IGλ clonotypes adjusted for off-target coverage |
|  | TCR richness | Measured based on the number of distinct TCR clonotypes adjusted for off-target coverage |
| | TCR $\alpha$ richness | Measured based on the number of distinct TCRα clonotypes adjusted for off-target coverage |
| | TCR $\beta$ richness | Measured based on the number of distinct TCRβ clonotypes adjusted for off-target coverage |
| | TCR $\delta$ richness | Measured based on the number of distinct TCRδ clonotypes adjusted for off-target coverage |
| | TCR $\gamma$ richness | Measured based on the number of distinct TCRγ clonotypes adjusted for off-target coverage |
| | IGH Shannon | Measured based on the number of distinct IGH clonotypes and their relative abundances.<br>$Shannon\ diversity = - \sum_{i=1}^n \frac{x_i}{N} \ln(\frac{x_i}{N}), x_i \neq 0; N = \sum_{i=1}^n x_i$ |
| | IG $\kappa$ Shannon | Measured based on the number of distinct IGκ clonotypes and their relative abundances.<br>$Shannon\ diversity = - \sum_{i=1}^n \frac{x_i}{N} \ln(\frac{x_i}{N}), x_i \neq 0; N = \sum_{i=1}^n x_i$ |
| | IG $\lambda$ Shannon | Measured based on the number of distinct IGλ clonotypes and their relative abundances.<br>$Shannon\ diversity = - \sum_{i=1}^n \frac{x_i}{N} \ln(\frac{x_i}{N}), x_i \neq 0; N = \sum_{i=1}^n x_i$ |
| | TCR $\alpha$ Shannon | Measured based on the number of distinct TCRα clonotypes and their relative abundances.<br>$Shannon\ diversity = - \sum_{i=1}^n \frac{x_i}{N} \ln(\frac{x_i}{N}), x_i \neq 0; N = \sum_{i=1}^n x_i$ |
| | TCR $\beta$ Shannon | Measured based on the number of distinct TCRβ clonotypes |
| | | and their relative abundances.<br>$Shannon\ diversity = - \sum_{i=1}^n \frac{x_i}{N} \ln(\frac{x_i}{N}), x_i \neq 0; N = \sum_{i=1}^n x_i$ |
| | TCR $\delta$ Shannon | Measured based on the number of distinct TCRδ clonotypes and their relative abundances.<br>$Shannon\ diversity = - \sum_{i=1}^n \frac{x_i}{N} \ln(\frac{x_i}{N}), x_i \neq 0; N = \sum_{i=1}^n x_i$ |
| | TCR $\gamma$ Shannon | Measured based on the number of distinct TCRγ clonotypes and their relative abundances.<br>$Shannon\ diversity = - \sum_{i=1}^n \frac{x_i}{N} \ln(\frac{x_i}{N}), x_i \neq 0; N = \sum_{i=1}^n x_i$ |

**Table S5.** Portion of zero values in the SBT features.

| SBT feature | OP-LUAD | OP-CRC | OP-BRCA | TCGA-RNA-Seq | TCGA-WXS |
| --- | --- | --- | --- | --- | --- |
| mtDNA copy number | 0.0% | 0.1% | 0.0% | - | 0.0% |
| 5S rDNA copy number | 0.0% | 0.0% | 0.0% | - | 0.0% |
| 18S rDNA copy number | 1.5% | 1.1% | 1.3% | - | 0.0% |
| 28S rDNA copy number | 0.1% | 0.2% | 0.0% | - | 0.0% |
| 45S rDNA copy number | 0.0% | 0.1% | 0.0% | - | 0.0% |
| fungal load | 0.0% | 0.0% | 0.0% | 0.0% | 0.0% |
| microbial load | 0.0% | 0.0% | 0.0% | 0.0% | 0.0% |
| protozoa load | 0.0% | 0.1% | 0.0% | 0.0% | 97.0% |
| viral load | 0.0% | 0.0% | 0.0% | 0.0% | 0.0% |
| IG $\kappa$ and $\lambda$ infiltration | 82.6% | 91.6% | 87.6% | 2.0% | 83.9% |
| IG $\kappa$ and $\lambda$ richness | 82.6% | 91.6% | 87.6% | 2.0% | 83.9% |
| BCR infiltration | 78.7% | 87.7% | 83.4% | 0.8% | 81.0% |
| IGH infiltration | 89.8% | 93.6% | 91.1% | 1.2% | 96.2% |
| IG $\kappa$ infiltration | 99.8% | 100.0% | 99.8% | 2.2% | 87.6% |
| IG $\lambda$ infiltration | 82.7% | 91.6% | 87.6% | 2.2% | 95.3% |
| T cell infiltration | 39.9% | 46.3% | 32.9% | 0.0% | 33.3% |
| T cell $\alpha$ infiltration | 39.9% | 46.5% | 32.9% | 0.0% | 53.0% |
| T cell $\beta$ infiltration | 62.7% | 67.5% | 57.5% | 15.9% | 59.1% |
| T cell $\delta$ infiltration | 99.6% | 99.6% | 99.1% | 12.9% | 97.3% |
| T cell $\gamma$ infiltration | 65.1% | 76.8% | 66.1% | 78.6% | 93.9% |
| BCR richness | 78.7% | 87.7% | 83.4% | 0.8% | 81.0% |
| IGH richness | 89.8% | 93.6% | 91.1% | 1.2% | 96.2% |
| IG $\kappa$ richness | 99.8% | 100.0% | 99.8% | 2.2% | 87.6% |
| IG $\lambda$ richness | 82.7% | 91.6% | 87.6% | 2.2% | 95.3% |
| TCR richness | 39.9% | 46.3% | 32.9% | 0.0% | 33.3% |
| TCR $\alpha$ richness | 39.9% | 46.5% | 32.9% | 0.0% | 53.0% |
| TCR $\beta$ richness | 62.7% | 67.5% | 57.5% | 15.9% | 59.1% |
| TCR $\delta$ richness | 99.6% | 99.6% | 99.1% | 12.9% | 97.3% |
| TCR $\gamma$ richness | 65.1% | 76.8% | 66.1% | 78.6% | 93.9% |
| IGH Shannon | 96.5% | 98.4% | 97.2% | 2.2% | 99.5% |
| IG $\kappa$ Shannon | 100.0% | 100.0% | 100.0% | 2.2% | 98.2% |
| IG $\lambda$ Shannon | 92.5% | 97.9% | 95.8% | 2.4% | 99.8% |
| TCR $\alpha$ Shannon | 47.9% | 56.4% | 41.2% | 0.0% | 79.6% |
| TCR $\beta$ Shannon | 91.6% | 93.6% | 86.5% | 31.0% | 86.9% |
| TCR $\delta$ Shannon | 100.0% | 100.0% | 99.9% | 29.0% | 99.8% |
| TCR $\gamma$ Shannon | 76.6% | 89.3% | 80.7% | 94.3% | 99.8% |

## Supplementary Figures

**Figure S1.**
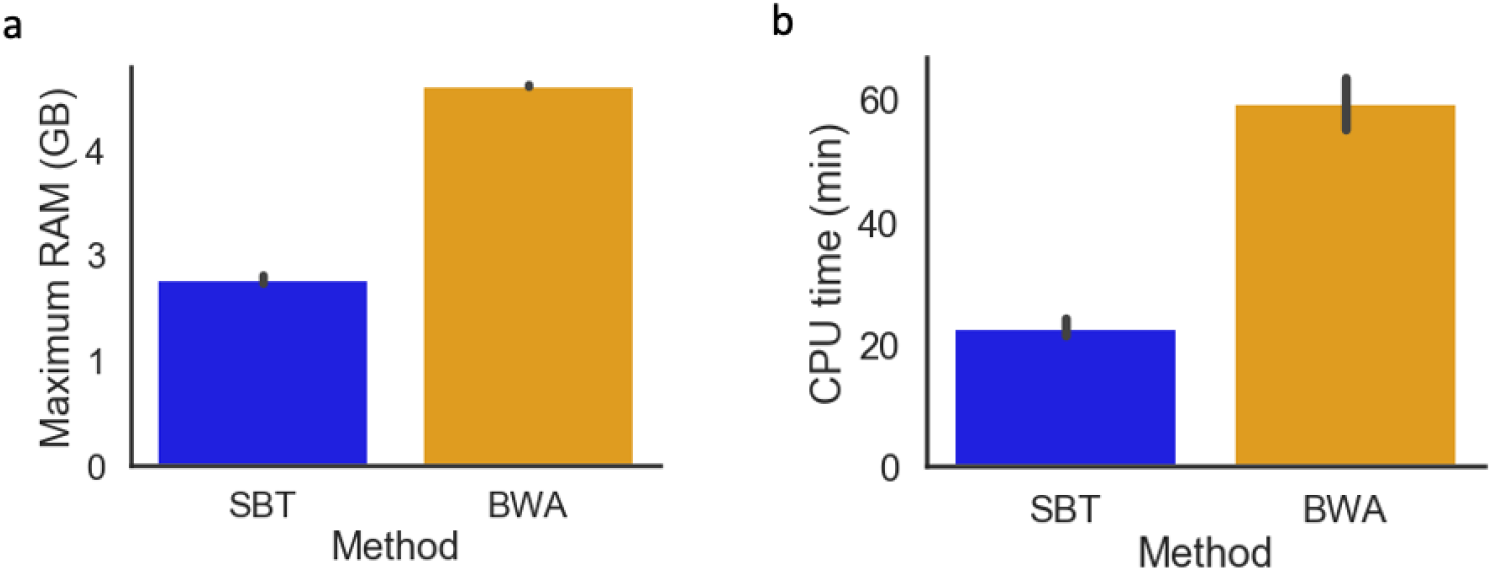
Barplot depicting (a) the maximum amount of RAM and b) the CPU time for running SBT and BWA methods. For each method, we report the mean value across 20 randomly selected samples from the OP-LUAD cohort.

**Figure S2.**
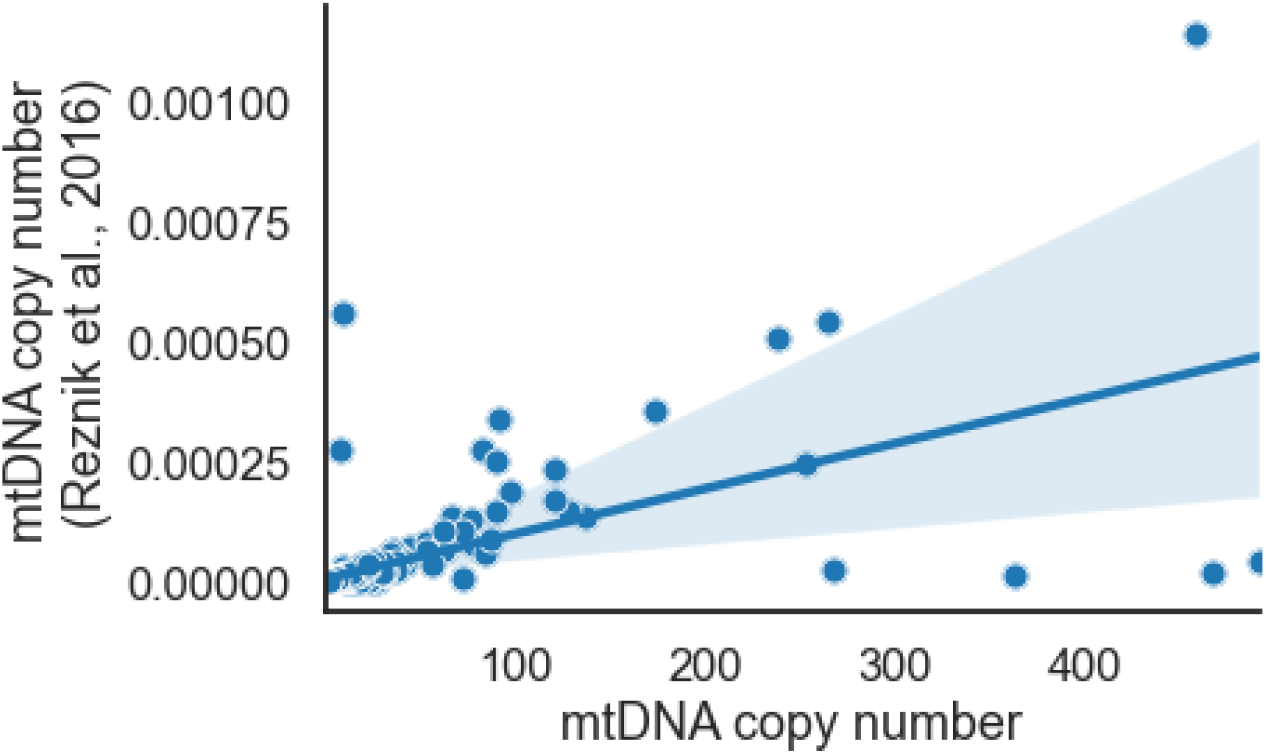
Correlation of WXS-based (TCGA LUAD) mtDNA dosage estimates obtained by SBT and by Resnik et al. ^11,12^ (Pearson correlation=0.61, p-value < 10^−20^).

**Figure S3.**
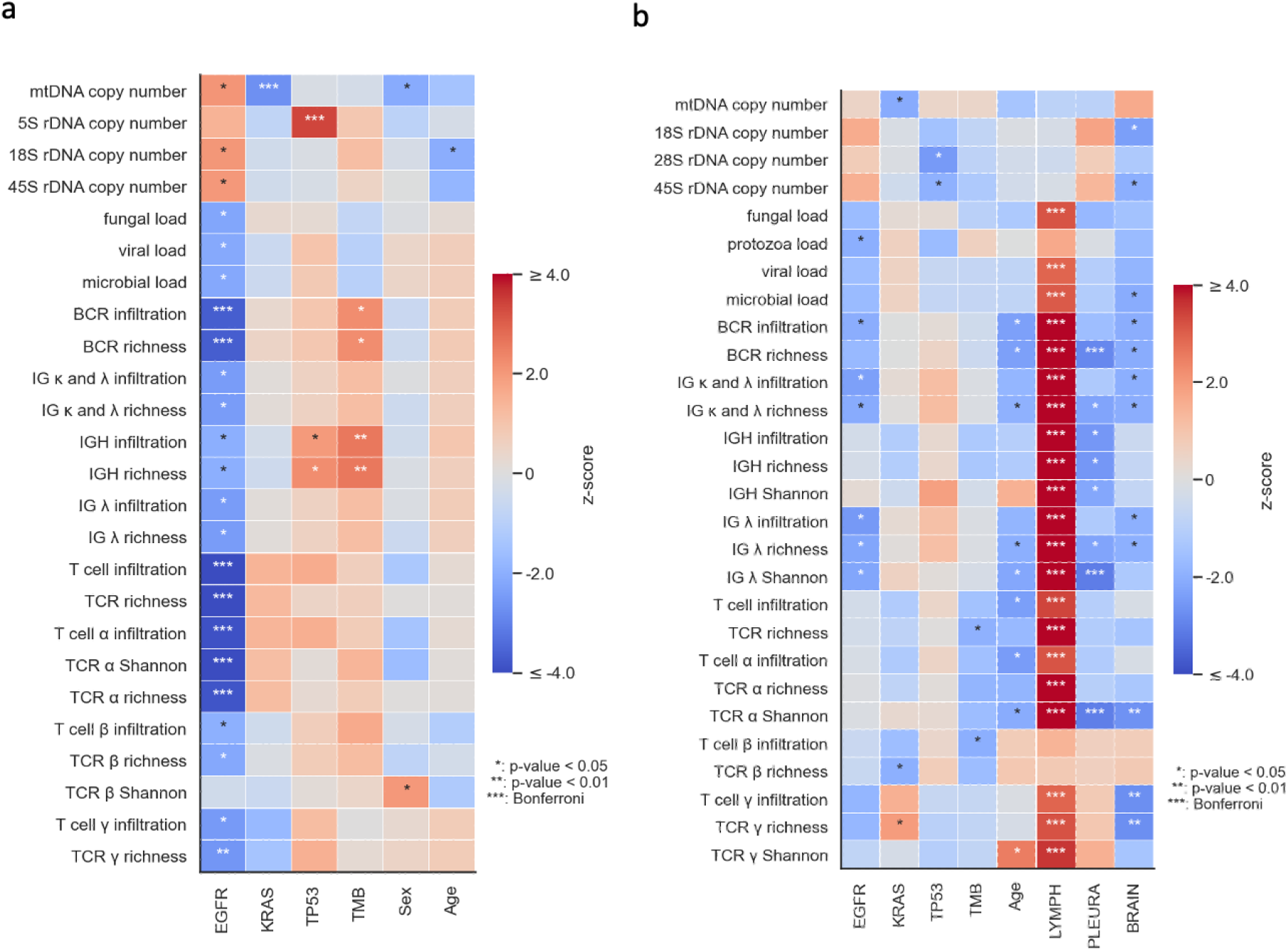
Association of SBT features with clinical factors and somatic mutations for (a) 1,352 LUAD samples biopsied from the LUNG and (b) 798 LUAD samples biopsied from non-LUNG sites, both datasets from the OP-LUAD cohort. Covariates are average off-target coverage, panel version, and tumor purity. Significant results are indicated with stars. Colors red and blue represent positive and negative association effect directions, respectively.

**Figure S4.**
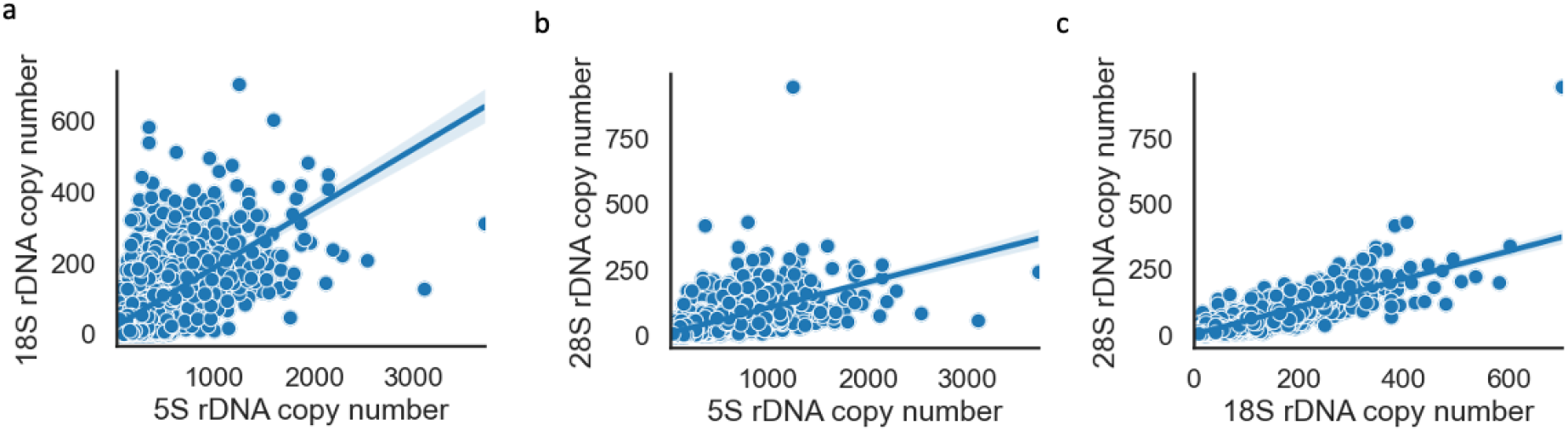
Correlation of rRNA copy number estimates across 2,920 samples from the OP-LUAD cohort: (a) Correlation of 18S-based (y-axis) and 5S-based rRNA copy number estimates (x-axis) (r=0.68, p-value<10^−20^); (b) Correlation of 28S-based (y-axis) and 5S-based rRNA copy number estimates (x-axis) (r=0.65, p-value<10^−20^); (c) Correlation of 28S-based (y-axis) and 18S-based rRNA copy number estimates (x-axis) (r=0.87, p-value<10^−20^).

**Figure S5.**
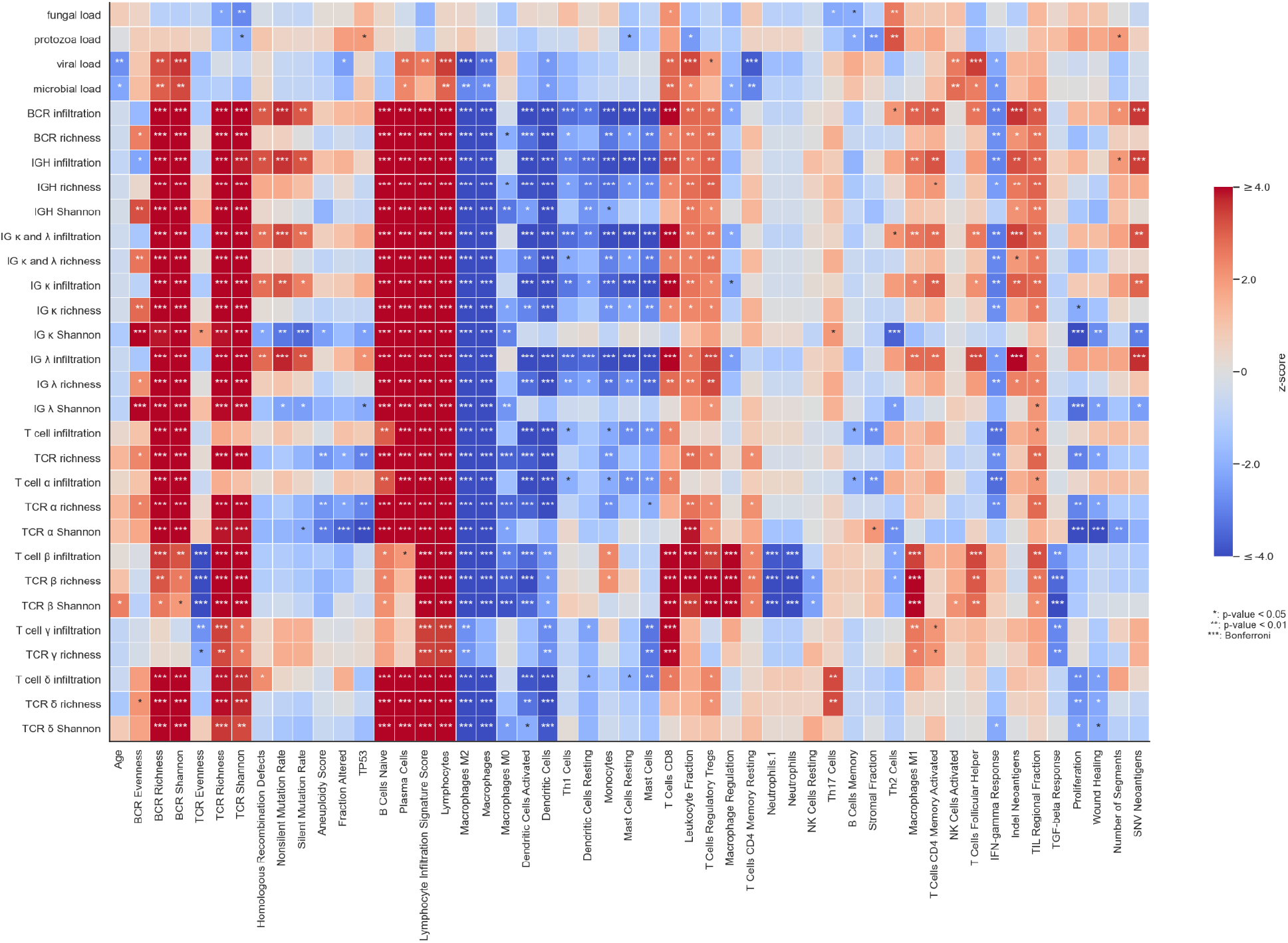
Association of SBT features with clinical factors and measurements from the immune landscape^15^ for TCGA-LUAD RNA-Seq samples. Covariates are the number of reads and tumor purity. Significant results are indicated with stars. Colors red and blue represent positive and negative association effect directions, respectively. We confirmed that SBT features inferred from RNA-seq were associated with immune landscape features (which were largely also inferred from RNA-seq in previous work). Given a large number of associations, we focused on broad patterns that were significant after the Bonferroni correction. Nearly all ImRep features were associated with increased measures of BCR/TCR expansion, T-Cells, B-Cells, Plasma Cells, and Lymphocyte infiltration; and associated with decreased measures of Macrophages, Dendritic Cells, Monocytes, and Proliferation (all Bonferroni p<0.05). In addition, IG features were broadly associated with mutation/neoantigen load; and TCR-related features were negatively associated with Neutrophil counts and proliferation.

**Figure S6.**
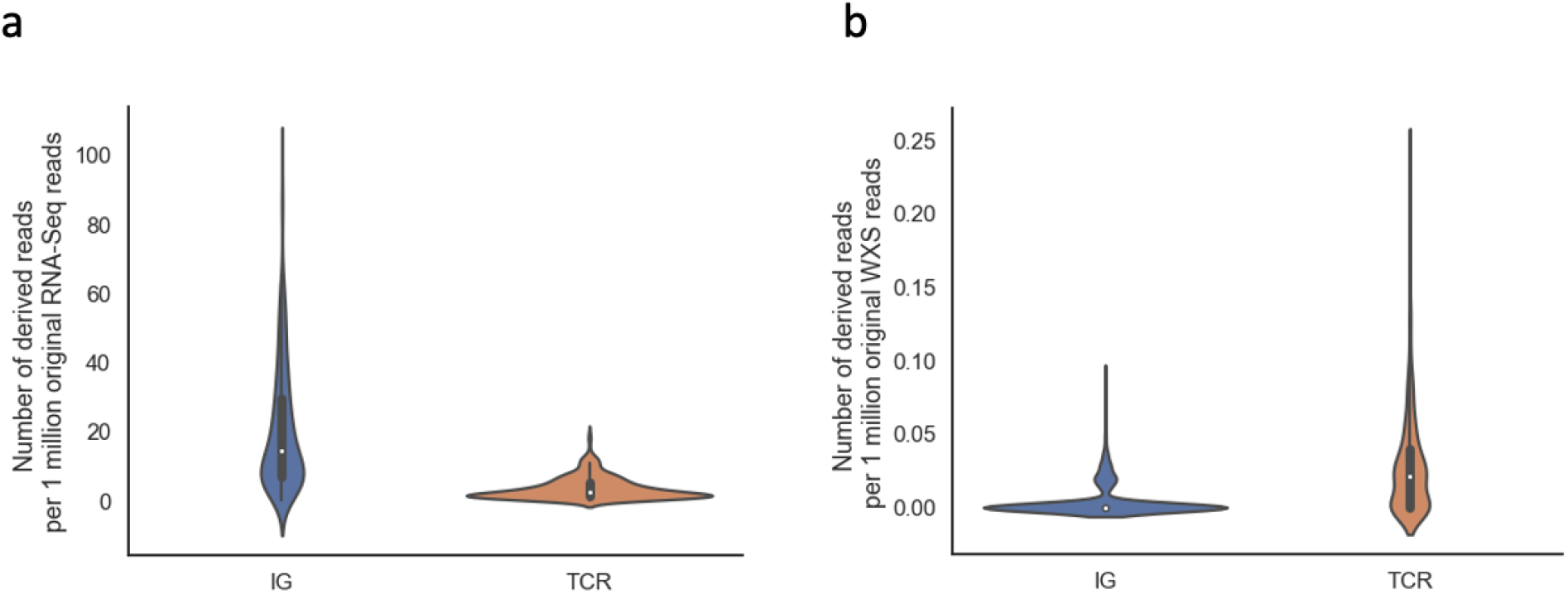
Comparison of receptor-derived reads provided by WXS and RNA-Seq (n=482) from TCGA-LUAD samples. a) Number of TCR/IG-derived reads per 1 million original RNA-Seq reads. b) Number of TCR/IG-derived reads per 1 million original WXS reads.

**Figure S7.**
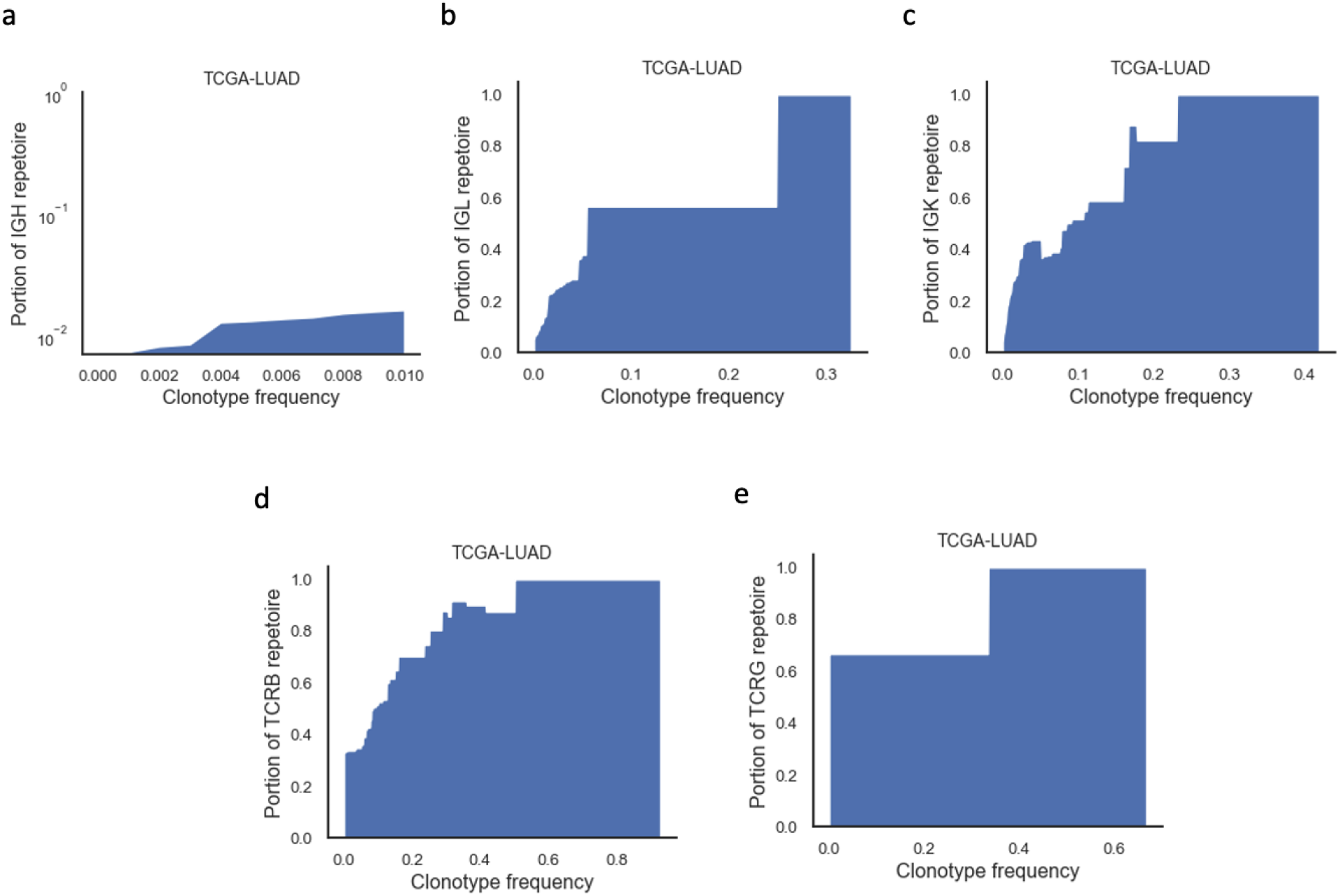
An area chart shows the proportion of the total (a) IGH, (b) IGL, (c) IGK, (d) TCRB, and (e) TCRG repertoire captured by ImRep on TCGA-LUAD depending on the minimum RNA-seq-confirmed clonotypes frequency considered. The x-axis corresponds to RNA-seq-confirmed clonotypes frequency *Z*. The y-axis corresponds to the fraction of assembled TCRB repertoire with clonotype abundances greater than *Z*. The total repertoire was defined as the sum of the RNA-seq-confirmed clonotypes abundances. Results on TCRA and TCRD were not presented, as few TCRA/TCRD-derived reads were detected from WXS samples.

**Figure S8.**
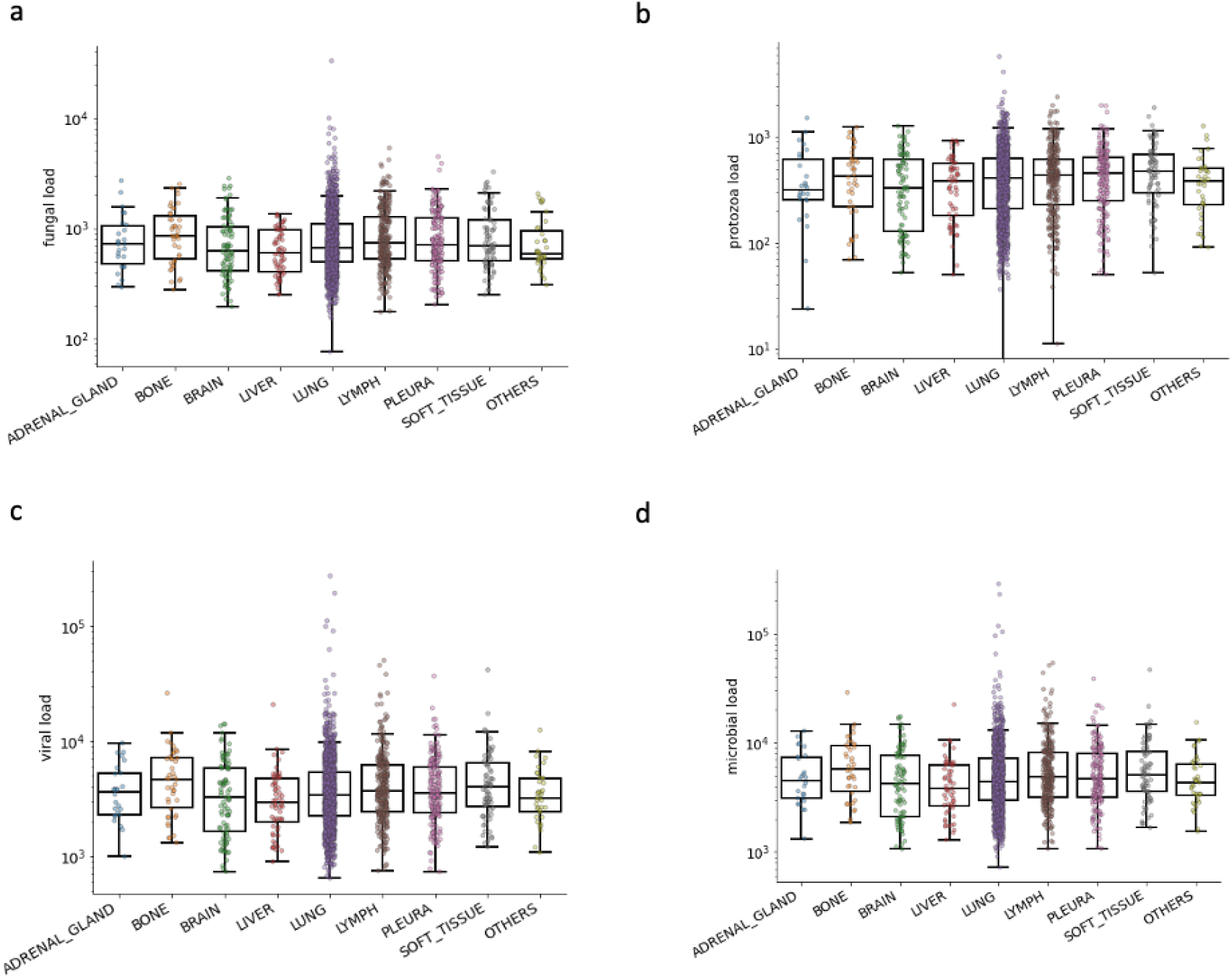
Comparison of (a) fungi load, (b) microbial load, (c) protozoa load, and (d) viral load across tissues of 2,150 samples from the OP-LUAD cohort. The distributions were compared using the Dunn-Bonferroni test, and all distributions are not statistically different (p-value>0.05).

